# Sequence alignment with *k*-bounded matching statistics

**DOI:** 10.1101/2025.05.19.654936

**Authors:** Tommi Mäklin, Jarno N. Alanko, Elena Biagi, Simon J. Puglisi

## Abstract

Finding high-quality local alignments between a query sequence and sequences contained in a large genomic database is a fundamental problem in computational genomics, at the core of thousands of biological analysis pipelines. Here, we describe a novel algorithm for approximate local alignment search based on the so-called *k*-bounded matching statistics of the query sequence with respect to an indexed database of sequences. We compute the *k*-bounded matching statistics, which capture the longest common suffix lengths of consecutive *k*-mer matches between query and target sequences, using the spectral Burrows-Wheeler transform, a data structure that enables computationally efficient queries. We show that our method is as fast and as accurate as state-of-the-art tools in several bacterial genomics tasks. Our method is available as a set of three kbo Rust packages that provide a command-line interface, a graphical user interface that runs in a browser without server-side processing, and a core library that can be accessed by other tools.

## Introduction

Genome analysis pipelines typically begin with the alignment of DNA sequences, in the form of, for example, sequencing reads or assemblies, against a known reference sequence. The resulting alignment describes compatibility between the two sequences and is the basis for many downstream analyses. In tasks where accuracy is essential and resources plentiful, the gold standard for alignment is to map short-read sequencing data against a reference genome assembled from both short and long reads [24]. However, particularly in microbial genomics, the largest current collections contain data from hundreds of thousands to millions of samples [18, 19, 14], making storage, distribution, and processing of large sequencing read sets the domain of only the largest institutions. This has led to the adoption of algorithms that perform essential tasks using only assembly data instead of sequencing reads, as specialized tools compress assembled genomes efficiently [9, 6] and tools operating on assemblies require less computational resources.

Tools for assembly comparison typically report the coordinates of aligned segments in a query relative to some reference, infer a consensus sequence consisting of nucleotide characters and gaps, and find insertions and deletions in the query. Accuracy in these three tasks depends highly on the quality of the input sequences [25]. Different tools tend to produce similar results with high-quality data, and thus focus shifted towards the development of new, faster, and more specialized algorithms and pipelines. In this paper, we devise and implement a new algorithm for solving the aforementioned comparison tasks by computing *k*-bounded matching statistics between query and reference sequences. For each position in the query sequence, a matching statistic describes the length of the sequence of characters preceding the position that are the same in both query and reference. The match length is capped at *k*characters, and so we use the term *k*-bounded matching statistics. We provide a formal definition in the Methods section.

Our approach, called kbo, is built on the recent spectral Burrows-Wheeler transform (SBWT) data structure, which allows rapid *k*-mer lookups in compact space [3]. These lookups can be extended to compute the *k*-bounded matching statistics by adding a longest common suffix array to the SBWT [1, 2]. We show that combining SBWT lookups with the *k*-bounded matching statistics information and a suffix match derandomization procedure enables retrieving the coordinates of matching regions in a query sequence even though the SBWT does not conserve the reference sequence location in its construction. Instead of storing the coordinates in the index, we stream the sequence where coordinate information should be preserved and index the sequence where location does not matter. As a whole, our algorithm represents a significant advance in *k*-mer matching methods, which have previously entirely ignored or discarded coordinate information and locality in favour of scalable lookups [4, 11].

In kbo we implement three main modes of operation, targeted at three different problems, that make use of *k*-mer lookups and *k*-bounded matching statistics. The first algorithm, *call*, is a specialized variant calling algorithm that reports insertions, deletions, and single-nucleotide variants in a query against some reference, describing local variations between two typically closely related sequences. Our second algorithm, *find*, compares a query and a reference and reports locally aligned segments in the query with an option to allow a small number of gaps and mismatches. Our last algorithm, *map*, performs reference-based alignment and produces a consensus sequence relative to the reference. The outputs from our three algorithms are useful in a number of bacterial genomics downstream analyses, ranging from genome-wide association studies to gene finding to phylogenetic inference, and their computational requirements are competitive with other state-of-the-art tools.

Beyond algorithmic novelty, the main advantage of kbo lies in its effective use of the SBWT data structure to perform comparisons in a rapid and parallelisable manner. We demonstrate that our implementation either matches or beats state-of-the-art methods in both accuracy and speed while retaining reasonable space requirements. Additionally, kbo requires no temporary disk space, making it ideal for massively parallel analyses. These low computational requirements also enable a fully in-browser implementation of our algorithms, greatly improving accessibility for non-experts and researchers with limited technical proficiency or resources. To our knowledge, kbo is the first fully-featured sequence aligner that can run entirely in the browser while also providing an efficient command-line client and a stable core library for use in other programs. We expect kbo to facilitate further democratization of bioinformatics and find adoption in fields processing large assembly collections.

## Methods

### Preliminaries

kbo is implemented on top of state-of-the-art string processing data structures and algorithms. In this section, we give an overview of the machinery we use to compute the *k*-bounded matching statistics. We provide only the key definitions necessary to be able to talk about the data structures on a technical level; the motivation and theory behind the definitions are described in detail in prior work [1, 2, 3].

The main workhorse of kbo is the spectral Burrows-Wheeler transform (SBWT) data structure. Roughly speaking, the SBWT lists the sets of outgoing edge labels from nodes in the de Bruijn graph of the input *k*-mers, such that these sets are sorted by the *colexicographic* (right-to-left lexicographic) order of the *k*-mers on the nodes.

To give the precise definition, we need to introduce the concepts of *k-spectrum* and *padded k-spectrum*.

#### Definition 1

(*k*-Spectrum) The *k*-spectrum 𝒮_*k*_(*S*) of string *S* is the set of distinct *k*-mers of *S*: {*S*[*i*..*i*+*k*−1] | *i* = 1, … , |*S*|−*k*+1}. The *k*-spectrum 𝒮_*k*_(*S*_1_, … , *S*_*m*_) of a set of strings *S*_1_, … , *S*_*m*_ is the union 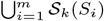.

Informally, a *padded k-spectrum* adds a minimal set of $-padded *dummy k*-mers to ensure that in the de Bruijn graph of 𝒮_*k*_(*S*_1_, … , *S*_*m*_) and the dummy *k*-mers, every non-dummy *k*-mer has an incoming path of length at least *k*.

#### Definition 2

(Padded *k*-Spectrum) Let *R* = 𝒮_*k*_(*S*_1_, … , *S*_*m*_) be a *k*-spectrum with alphabet Σ, and let *R*^′^ ⊆ *R* be the set of *k*-mers *Y* such that *Y* [1..*k*− 1] is not a suffix of any *k*-mer in *R*. The padded *k*-spectrum is the set 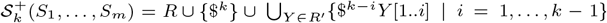, with special character $∉ Σ and $ *< c* for all *c* ∈ Σ.

For example, if *S*_1_ = ACGT, *S*_2_ = GACG and *k*= 3, then 𝒮_3_(*X*_1_, *X*_2_) = {ACG, CGT, GAC} and 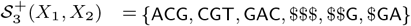.

We are now ready to define the SBWT.

#### Definition 3

(Spectral Burrows-Wheeler transform (SBWT) [3]) Let *R*^+^ be a padded *k*-spectrum and let *S*_1_, … , *S*_|*R*|_ be the elements of *R*^+^ in colexicographic order. The SBWT is the sequence of sets *A*_1_, … , *A*_|*R*|_ with *A*_*i*_ ⊆ Σ such that *A*_*i*_ = ∅ if *i >* 1 and *S*_*i*_[2..*k*] = *S*_*i*−1_[2..*k*], otherwise *A*_*i*_ = {*c* ∈ Σ | *S*_*i*_[2..*k*]*c* ∈ *R*^+^}.

We refer the reader to [1, 2, 3] for examples. The Longest Common Suffix array is the key to efficient computation of the *k*-bounded matching statistics (to be defined later in this section):

#### Definition 4

(Longest Common Suffix (*LCS*) Array [1]) Let *R*^+^ be a padded *k*-spectrum and let *X*_*i*_ be the colexicographically *i*-th *k*-mer of *R*^+^. The *LCS* array is an array of length |*R*^+^| s.t. *LCS*[1] = 0, and for *i >* 1, *LCS*[*i*] is the length of the longest common suffix of *S*_*i*−1_ and *S*_*i*_.

In the above definition, we consider the empty string as a common suffix of any two *k*-mers, so that the longest common suffix is well-defined for any pair of *k*-mers. An example *LCS* array is shown in Figure 1.

**Fig. 1:**
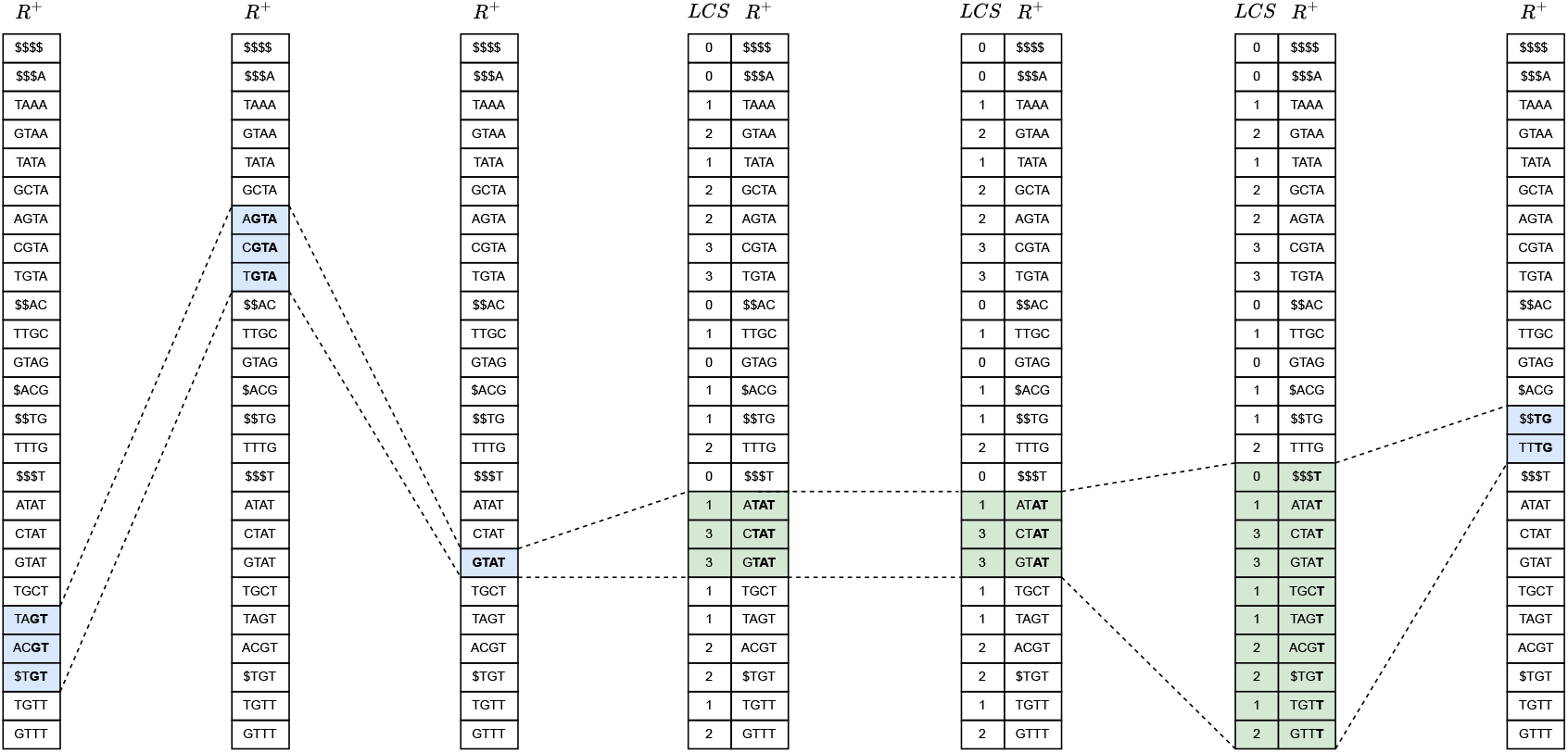
Illustration of *ExtendRight* (resulting in blue intervals) and *ContractLeft* operations (resulting in green intervals) to match the string GTATG with *k*= 4 against a reference S = {TGTTTG, TTGCTAT, ACGTAGTATAT, TGTAAA}. This match produces the matching statistics vector *MS* = {1, 2, 3, 4, 2} and derandomizes to the vector *DMS* = {1, 2, 3, 4, 0} if the threshold used is *t* = 2.

When the *k*-mers in *R*^+^ are sorted colexicographically, the subset of *k*-mers in *R*^+^ that share a string *α* as a suffix are next to each other, forming a contiguous range. This is called the *colexicographic range* of *α*. The colexicographic range of a string *α* that is *longer* than *k*is defined as the range of *k*-mers in the sorted list of *R*^+^ that have the last *k*characters of *α* as a suffix, making the range either singleton or empty. The SBWT provides two operations on colexicographic ranges [*l, r*] of any string *α*:

- ExtendRight(*l, r, c*) returns the colexicoraphic range of string *αc* (possibly an empty range).
- ContractLeft(*l, r*), with [*l, r*] being the colexicographic range of *α*, returns the colexicographic range of *α*[2..|*α*|].

See Alanko et al. [2] for details on how to implement these operations using the SBWT. Figure 1 provides an illustration of both ExtendRight and ContractLeft operations.

The *k*-bounded matching statistics are defined as follows:

#### Definition 5

(*k*-bounded matching statistics [2]) The *k*-bounded matching statistics for a query string *Q* against a set of reference strings *S*_1_, … , *S*_*m*_ is a vector *MS*_*k*_[1..|*Q*|] such that *MS*_*k*_[*i*] is the largest integer *l* ≤ *k*such that *Q*[*i* − *l* + 1..*i*] is a substring of at least one reference *S*_1_, … , *S*_*m*_.

In what follows, we may drop the subscript of *MS*_*k*_ since *k*is assumed to be fixed at indexing time.

The padded *k*-spectrum *R*^+^ provides an equivalent formulation that is easier to compute: the value of *MS*[*i*] is the length *l* of the longest *suffix of a k-mer* in the padded spectrum of *T*_1_, … , *T*_*m*_ that matches *Q*[*i* − *l* + 1..*i*]. The *k*-mers in *R*^+^ that have suffix *Q*[*i* − *l* + 1..*i*] are adjacent in the colexicographically sorted list of *k*-mers in *R*^+^ (the colexicographic range of *Q*[*i* − *l* + 1..*i*]). Algorithm 1 shows pseudocode to compute the MS vector and the colexicographic ranges [*l, r*] corresponding to the longest match at each position.

### Identifying statistically significant match positions

For each query *Q*, we use Algorithm 1 to compute the matching statistics vector *MS*[1..|*Q*|]. The matching statistic *MS*[*i*] is the length of the longest match ending at position *i* in *Q*, up to length *k*(See Definition 5). Even a random query will likely match some number of nucleotides at every position by chance alone. Our goal in this section is to derive a threshold for a statistically significant match length.

This threshold must be a function of the number of *k*-mers in the index: the larger the amount of indexed data, the longer matches we expect to find by chance. Let us denote the set of *k*-mers in the index with ℐ. We use a model where all the *k*-mers in ℐ are independent random strings. The match length distribution at a given position against a single random *k*-mer follows a geometric distribution truncated at *k*. To simplify the formulas, we assume an untruncated geometric distribution. This is a reasonable approximation since for values *k*≥ 30 that we use, the tail probability of random suffix match lengths close to *k*is vanishingly small. With these assumptions, if *X* is the random variable denoting the length of the match against a single *k*-mer, the cumulative distribution *P* (*X* ≤ *t*) is given by:

#### Algorithm 1

*k*-bounded matching statistics.

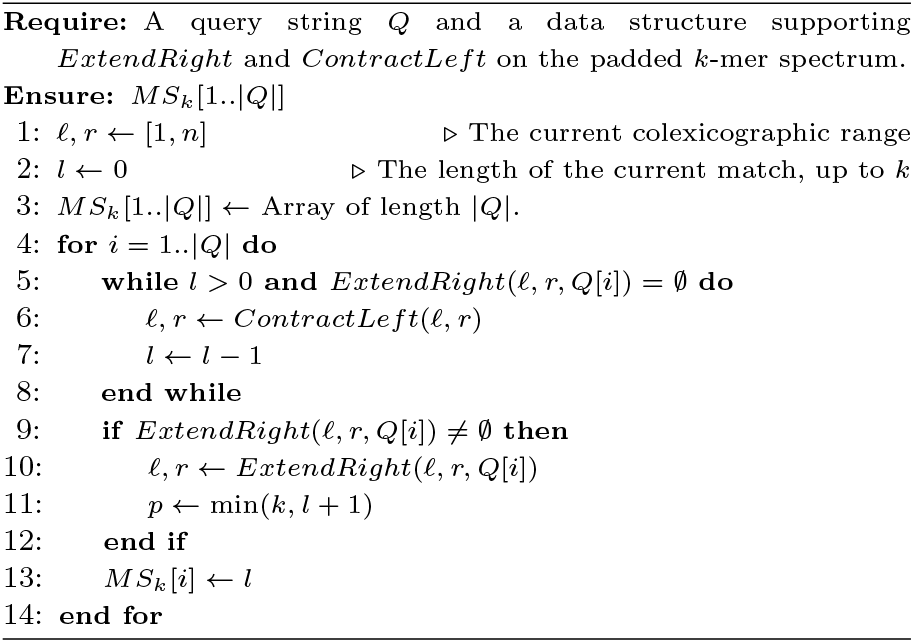

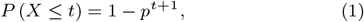

where *p* is the probability that two nucleotides match by chance, e.g., *p* = 1*/*4. Now, let *M* be a random variable denoting the length of the longest suffix match ending at some position *Q*[*i*] against the entire index ℐ containing *n* distinct *k*-mers. Since the *k*-mers in ℐ are assumed to be independent, the CDF of the maximum match length is:

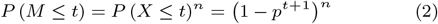

Suppose we want to consider a match as statistically significant if it occurs with probability at most *r* in our model. We can determine the corresponding match length *t*_*p*_ by setting the CDF in Equation 2 to greater than 1 − *r* and solving for *t*. We obtain:

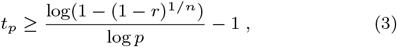

For example, for a small genome with *n* = 10^6^ distinct *k*-mers, nucleotide match probability *p* = 0.25 and false positive tolerance *r* = 10^−6^, we have *t*_*p*_ ≈ 18.9.

### Derandomizing *k*-bounded matching statistic vectors

Using the threshold *t*_*p*_ from Equation (3), we *derandomize MS* by replacing match lengths below the threshold *t*_*p*_ with values derived from nearby values. A key fact to understand about the *MS* vector is that *MS*[*i*] is always at most *MS*[*i* − 1] + 1. For typical data in practice, the general shape of the *MS* vector is a mountainscape with slow climbs and sharp falls, interspersed with with noisy valleys below the significant match threshold (see Figure 2). The idea of derandomization is to extrapolate the climbs backward to eliminate the noisy segments. The process is detailed in Algorithm 2. After this processing, we can easily read off the lengths of gaps in the matches, as shown in the next section.

**Fig. 2:**
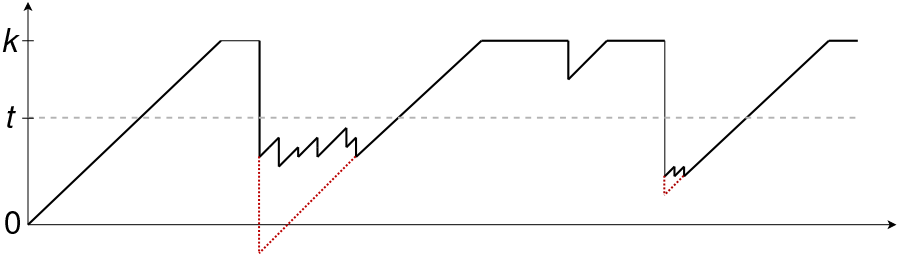
Derandomizing the MS vector. The vertical axis is the length of a match (up to the *k*-mer size *k*) ending at a given position in the query (horizontal axis). The black line shows values of the *k*-bounded matching statistics. The dashed grey horizontal line marks the noise threshold *t*. The dotted red line shows the derandomized values.

#### Algorithm 2

Derandomizing the MS vector

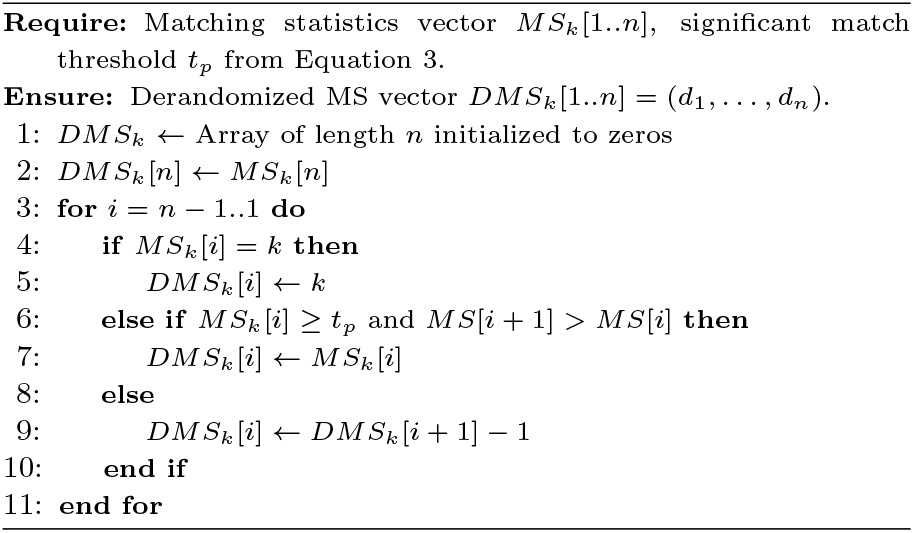

### Translating derandomized matching statistics into alignment events

The derandomized MS vector can be translated into a character representation of the compatibility between the query and a reference sequence. For example, a single nucleotide substitution or insertion at position *i* will show up as run from 0 to *k*in *DMS*_*k*_[*i*..*i* + *k*]. These can be distinguished from deletions and recombinations, which exhibit different kinds of characteristic patterns. All the possible cases are displayed in Figure 3. We represent character compatibility events using the following set of symbols:

**Fig. 3:**
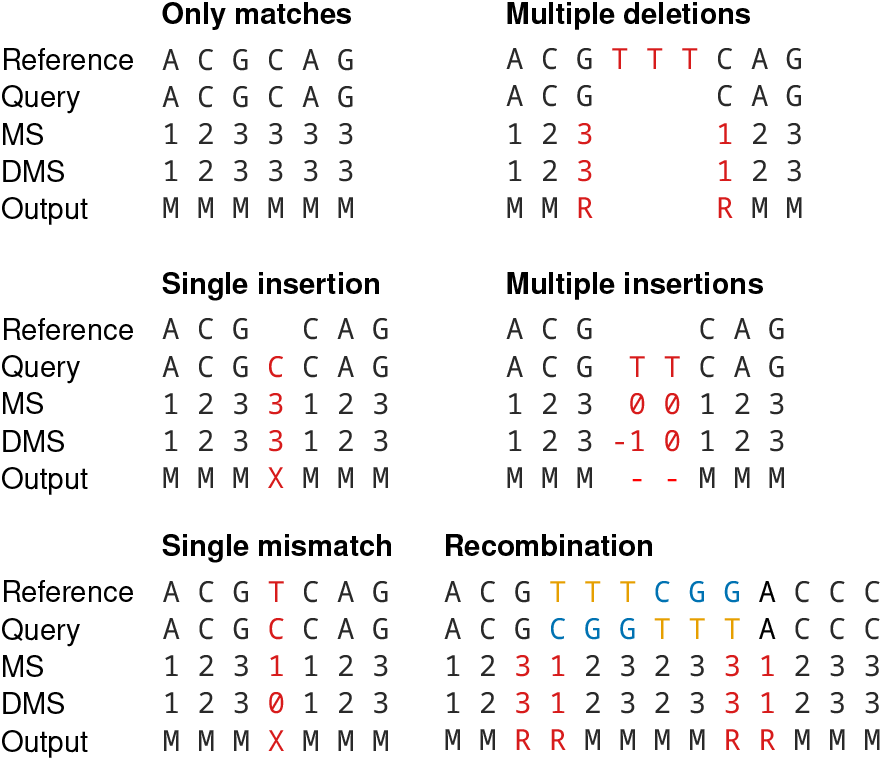
Illustration of the results of applying the translation procedure to a derandomized matching statistics vector. Examples in both panels assume *k*-mer size 3 and derandomization threshold *t* = 2. MS is short for matching statistics and DMS for derandomized matching statistics.

`M` : Match between query and reference.

**-** : Bases in the query that are not found in the reference.

`X` : Single base mismatch *or* single base insertion into the query.

`R` : Two consecutive `R`s signify a discontinuity in the alignment. The right R is at the start of a *k*-mer that is not adjacent to the last character in the *k*-mer corresponding to the left `R`. This implies either a deletion of unknown length in the query, or an insertion of *k*-mers from elsewhere in the reference into the query.

Algorithm 3 shows how these events can be detected with a single pass over the *DMS*_*k*_ array. Figure 3 shows examples of the distinct cases. Note how these examples illustrate that the procedure is not able to distinguish between a single base insertion and a single base substitution. This problem can be resolved by adding a *refinement* step to our translation, which resolves this conundrum by using the nucleotide sequence of the reference.

#### Algorithm 3

Translating the DMS vector

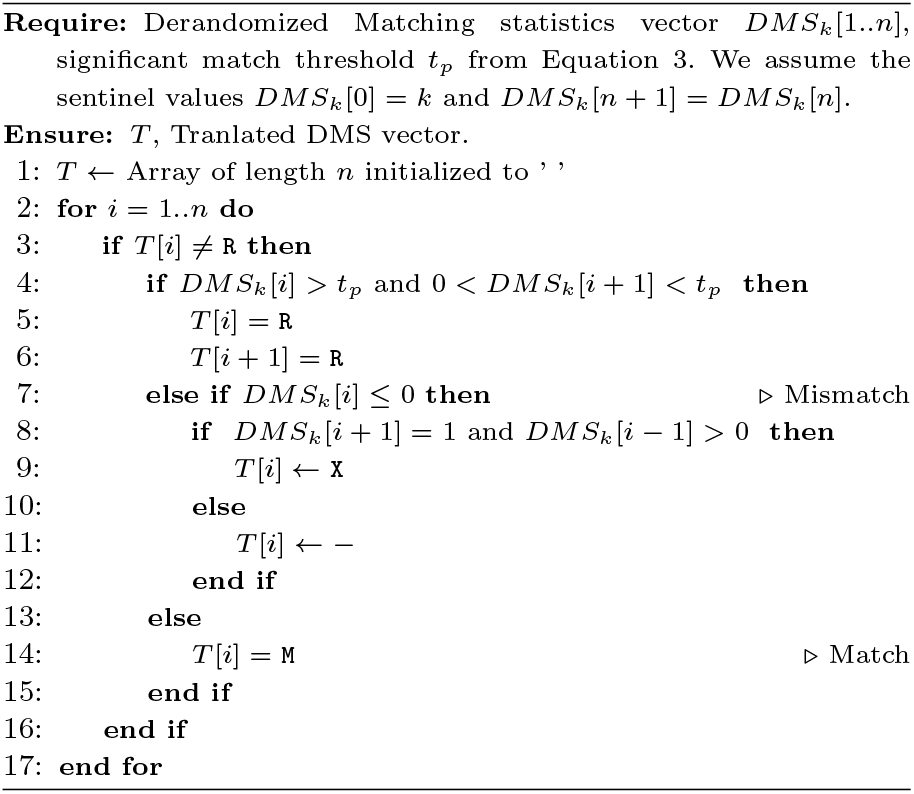

### Variant calling using matching statistics

kbo also implements a variant calling algorithm to detect and report all short variants between a pair of sequences: a reference *R* and a query *Q*. A variant is defined as a triple (*i, v*_*r*_, *v*_*q*_), which means that at location *R*[*i*], the string *v*_*r*_ has been replaced with string *v*_*q*_. This framework encompasses all types of short variations: If *v*_*r*_ is empty, the variant is an insertion, and if *v*_*q*_ is empty, the variant is a deletion. If |*v*_*r*_| = |*v*_*q*_| = 1, we have a single nucleotide substitution. If the lengths of |*v*_*r*_| *>* 1, |*v*_*q*_| *>* 1 and |*v*_*r*_| ≠ |*v*_*q*_| the variant affects multiple bases and includes substitutions and indels.

To find variants between the sequences *R* and *Q*, we first build the SBWT indices of both *R* and *Q*, and compute the *k*-bounded matching statistics array *MS*[1..|*Q*|] of *Q* against *R*. The idea is to analyze positions where the match length drops from significant to insignificant. A position *i* is classified as a *significant drop* iff *MS*[*i* − 1] ≥ *t*_*p*_ and *MS*[*i*] *< t*_*p*_, where *t*_*p*_ is the significant match threshold derived in Section “Identifying statistically significant match positions”. The position *i* − 1 just before the drop is called a *significant peak*.

Each significant peak position *i* is analyzed by finding the shortest unique significant match nearest to the right of *i*, that is, the smallest *j > i* such that *MS*[*j*] ≥ *t*_*p*_ and the longest match ending at *Q*[*j*] is a suffix of only one *k*-mer of *R*. This unique match is used as an anchor to resolve the potential variant between *R* and *Q*. This is done by comparing the *k*-mer *α* = *Q*[*j* − *k*+ 1..*j*] and the unique *k*-mer *β*, in the SBWT of *R*, that has suffix equal to *Q*[*i* + 1..*j*].

Denoting the length of the longest common suffix of *α* and *β* with *m*, the potential variant ends just before position *e* = *k*−*m*+1 in both *α* and *β*. To find the variant sequences *v*_*r*_ and *v*_*q*_, we compute the matching statistics *MS*^*α*^[1..|*α*|] of *α* against *R* and *MS*^*β*^[1..|*β*|] of *β* against *Q*. The starting points *s*_*α*_ in *α* and *s*_*β*_ in *β* are at the rightmost (if they exist) significant drops of *MS*^*α*^ and *MS*^*β*^, respectively. That is, *v*_*q*_ = *α*[*s*_*α*_..*e*) and *v*_*r*_ = *β*[*s*_*β*_..*e*). However, we must be careful because in case of deletions and insertions, the predicted starting point may fall *after* the predicted ending point (see Fig. 4, Case 3), or *s* and *e* may coincide (Case 2). We define gap(*α*) = *e* − *s*_*α*_ and gap(*β*) = *e* − *s*_*β*_ (See Fig. 4). If one or both of these gaps is negative or 0, we have an indel. If gap(*α*) *>* gap(*β*), we have an insertion in *Q*. On the other hand, if gap(*α*) *<* gap(*β*), we observe a deletion in *Q*.

**Fig. 4:**
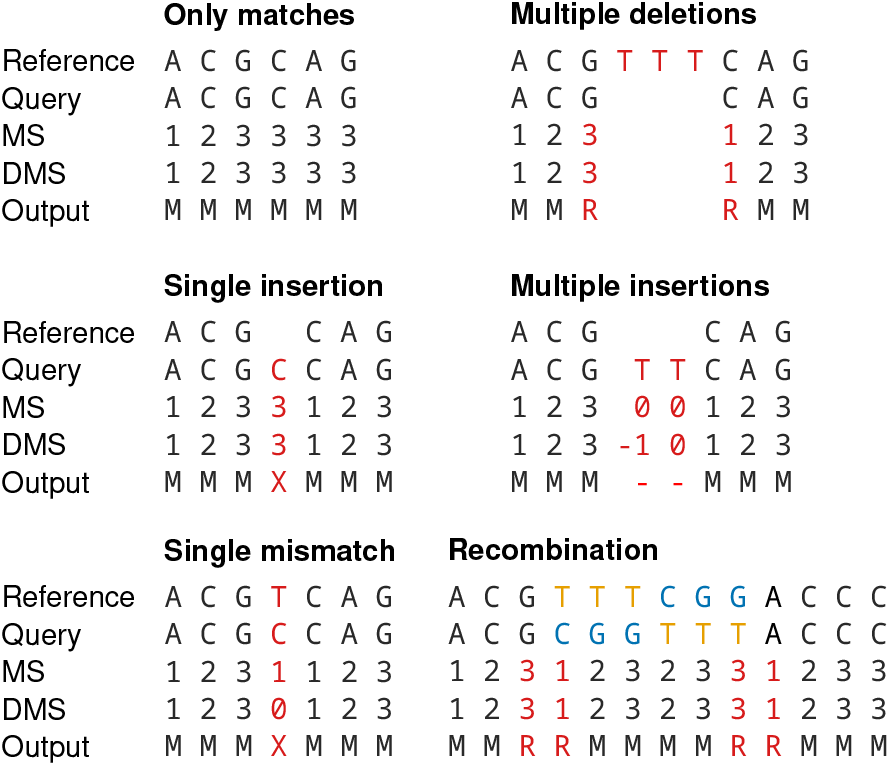
Examples of variant calling outputs with *k*= 9. The first case is the most basic one: We have a substitution of CTC with AG, which are found in the range [*s*..*e*) in each sequence. In the second case, with have an insertion of one character which is not equal to the preceding character in the query, making one of the intervals [*s*..*e*) empty, indicating an indel. In the third case, we have an insertion of characters that are equal to the preceding character. In this case, the predicted start and end points cross, making the gaps negative, but the difference in the length of the gaps still gives the length of the insertion.

### Filling gaps in the translated alignment

Translated output from kbo may contain regions consisting of single or consecutive ‘-’ characters, which we call gaps, that are false negatives. These are primarily caused by multiple changes occurring within an interval shorter than the *k*-mer size, which leads the derandomization algorithm to mark the entire interval as a gap as the matching statistics within the region never exceeds the threshold. If these gaps are short enough that a single *k*-mer overlaps the entire region, variant calling will resolve the true sequence, but for longer gaps a dedicated gap filling algorithm, which we now describe, is required.

We start by defining the *nearest unique context* of a gap. The nearest unique context *ν* is the colexicographic range [*l, r*] of size 1 (*r* = *l*) of a single *k*-mer *γ*, from the SBWT of the reference, whose suffix matches the characters to the right of the gap. We require suffix matches of at least the derandomization threshold *t*_*p*_ in order to avoid spurious *k*-mers. This implies that *γ* always overlaps at least *t*_*p*_ characters to the right of the gap. The first step in our gap filling algorithm is to search for the nearest unique context and, if it does not exist, the gap is considered unfillable.

If *γ* exists, we attempt to find a sequence *λ* that extends *γ* to the left so that the prefix of *λ* matches the sequence to the left of the gap by *t*_*p*_ characters. We construct *λ* iteratively by prepending each possible character {`A, C, G, T`} in turn to the concatenated sequence *λ*_*i*−1_ · *γ*, where *λ*_0_ is an empty sequence and *λ*_*i*−1_ is the concatenated sequence from the previous iteration. At each iteration we check that 1) the first *k*characters in the extended sequence *λ*_*i*_ have a colexicographic range with size 1 in the SBWT, and 2) only one of the four possible extensions has a colexicographic range with size 1. If either of these conditions fails before the prefix of *λ*_*i*_ matches the sequence left of the gap by *t*_*p*_ characters, we say that a valid extension *λ* does not exist. Note that *γ* may already fulfil these criteria if the gap is shorter than *k*− 2*t*_*p*_, in which case the valid extension is the empty sequence *λ*_0_.

If a valid left extension *λ* exists, we extract the subsequence *δ* from the concatenated sequence *λ* · *γ* by taking the characters that overlap the gap, 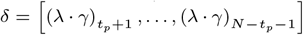 where *N* is the total size of *λ* · *γ*, and apply a series of checks to determine whether to use *δ* to fill the gap. The first check is that the size of *δ* must equal the size of the gap, meaning there are no insertions or deletions in *λ*. The second check is that *δ* and *λ* must meet at least one of the following three requirements: 1) *λ* is an empty sequence, or 2) all characters in *δ* except 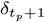 and 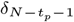 match the reference, or 3) the probability of generating all consecutive matches in *δ* is less than *t*_*p*_. We compute 3) using products of the CDF of maximum match lengths (Definition 2). If *δ* passes these checks, we replace the gap characters with the sequence *δ*.

### Implementation

We provide kbo in the form of three software packages written fully in the Rust programming language: kbo^1^ contains the implementations of our algorithms and is intended for usage in other programs, kbo-cli^2^ provides a command-line client, and kbo-gui^3^ is a Rust-to-WebAssembly graphical user interface that runs in a browser with no server-side processing or data transfer. Both the command-line and graphical user interfaces provide full access to all features of kbo. An overview of the operation of kbo is given in Figure 5.

**Fig. 5:**
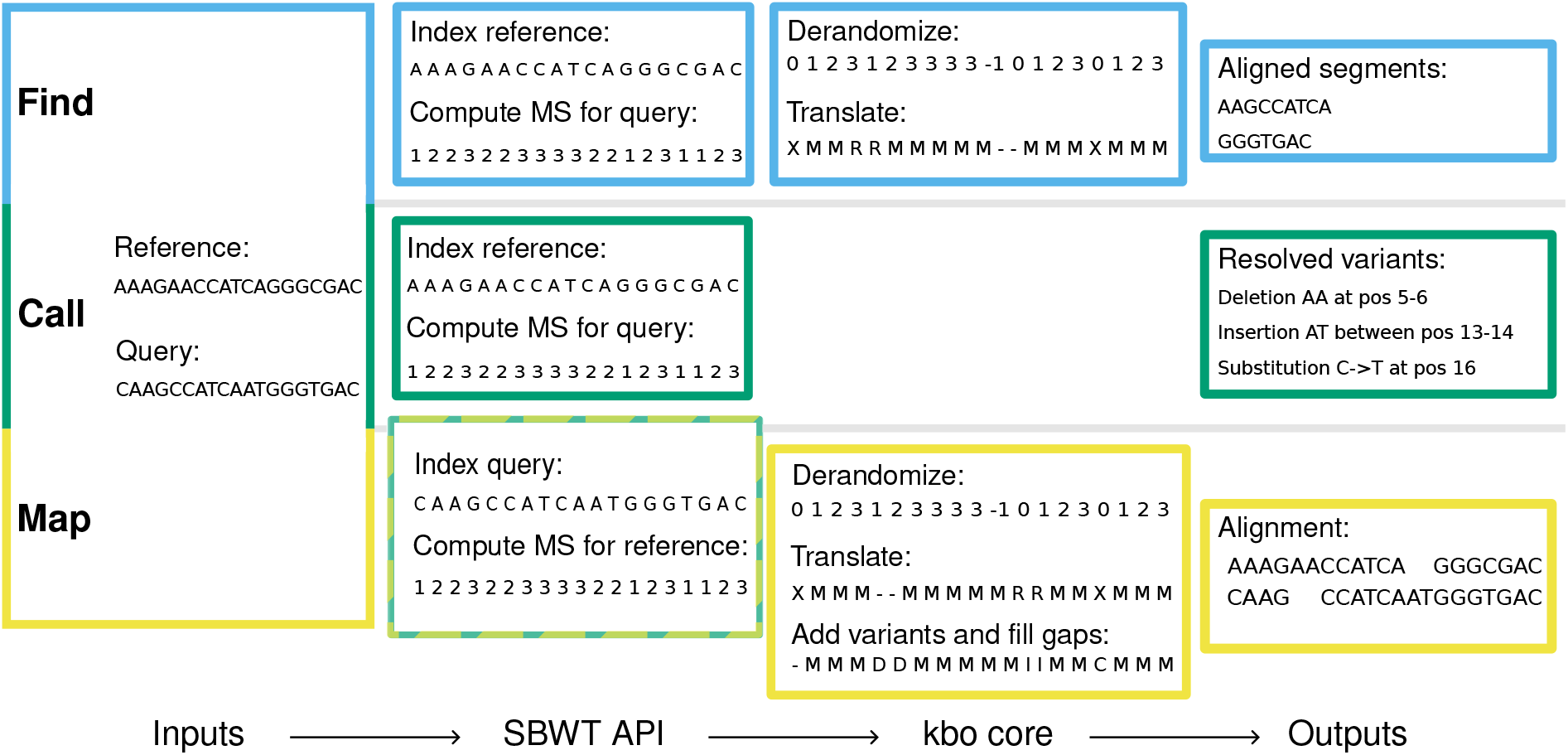
Overview of the three main algorithms in kbo: find, call, and map. The figure shows which parts of the SBWT and kbo core libraries each algorithm makes use of and illustrates example inputs and outputs to and from the user interface implementations. This figure was created with *k*-mer size 3 and derandomization threshold 2. In the translated output, ‘D’ denotes a deletion, ‘I’ an insertion, and ‘X’ a single nucleotide polymorphism. In the translated output and the Alignment box sequences, ‘-’ denotes base pairs that are not found in the reference.

## Use cases

We demonstrate kbo on three common bacterial genomics tasks and present results for both accuracy and compute resource usage. These demonstrations were conducted using the command-line interface (kbo-cli v0.2.0) running on a 2021 Dell XPS 13 laptop with an Intel i7-1165G7 4-core processor and 16GB RAM. All tools were run with the default parameters unless otherwise stated.

### Variant calling with ‘kbo call’

We benchmarked the variant calling algorithm in kbo by reproducing an experiment conducted in an earlier study [25]. Briefly, in that study, seven very closely related (less than 10 nucleotides difference) *Staphylococcus aureus* assemblies were created with different assemblers and sequencing technologies, including short-read and long-read sequencing of the same isolates. The study compared variant calls obtained from assemblies only against the gold standard read-based calls on a hybrid-assembled 2.9Mb long reference genome. We compare the variant calls from kbo against calls generated using multiple different approaches: 1) based on genome alignment (MUMmer4[16]), 2) a pipeline converting the assemblies into pseudo reads using wgsim, aligning them against the reference with BWA-MEM [15], and calling the variants with freebayes [12] (called Shred, because the pipeline shreds the assemblies into pseudo-reads using wgsim), and 3) a split *k*-mer based approach (SKA [10]). We used default parameters for all three tools and *k*= 51 and *t*_*p*_ = 10^−8^ for kbo.

Our results (Tables 1 and 2) show that, like MUMmer4 and Shred, kbo achieves perfect sensitivity and successfully calls all SNVs and indels that are present in the gold standard. When measured by precision, kbo is markedly better at reporting fewer false positives, achieving two times better precision in SNV calling and three times better in resolving indels. Supplementary Figure 1 shows how the results from kbo change when the *k*-mer size and the maximum error probability parameters are varied.

**Table 1.** Results from four tested methods in resolving single-nucleotide substitutions. The ground-truth was determined with read-based variant calling against a hybrid-assembled reference genome. Values are rounded to 2 most significant digits.

| Method | TP | FN | FP | Sensitivity | Precision |
| --- | --- | --- | --- | --- | --- |
| kbo | 782 | 0 | 1630 | 1.00 | 0.32 |
| MUMmer4 | 782 | 0 | 4669 | 1.00 | 0.14 |
| Shred | 782 | 0 | 4001 | 1.00 | 0.16 |
| SKA | 374 | 408 | 7431 | 0.48 | 0.05 |

**Table 2.**
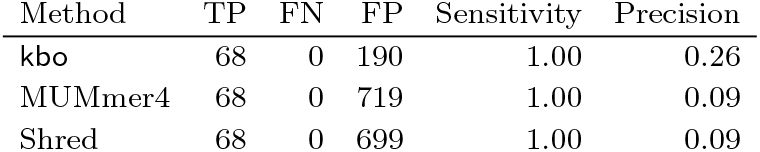
Results from three tested methods in resolving insertions. SKA does not resolve insertions and deletions so it is omitted. The ground-truth was determined with read-based variant calling against a hybrid-assembled reference genome. Values are rounded to 2 most significant digits.

| Method | TP | FN | FP | Sensitivity | Precision |
| --- | --- | --- | --- | --- | --- |
| kbo | 68 | 0 | 190 | 1.00 | 0.26 |
| MUMmer4 | 68 | 0 | 719 | 1.00 | 0.09 |
| Shred | 68 | 0 | 699 | 1.00 | 0.09 |

By resource usage kbo is the fastest method, taking an average of 1.6s seconds to process a single pairwise comparison (Fig. 6a). The peak memory usage of kbo is comparable to the other *k*-mer based method, SKA, requiring 401MB per comparison on average (Fig. 6b). Since kbo performs all operations fully in-memory, the tool is the only one of the four tested that requires no temporary disk space (Fig. 6c). Overall the resource usage of kbo is mainly determined by the *k*-mer size parameter. Supplementary Figure 2 shows how the runtime and memory usage change with different values of *k*.

**Fig. 6:**
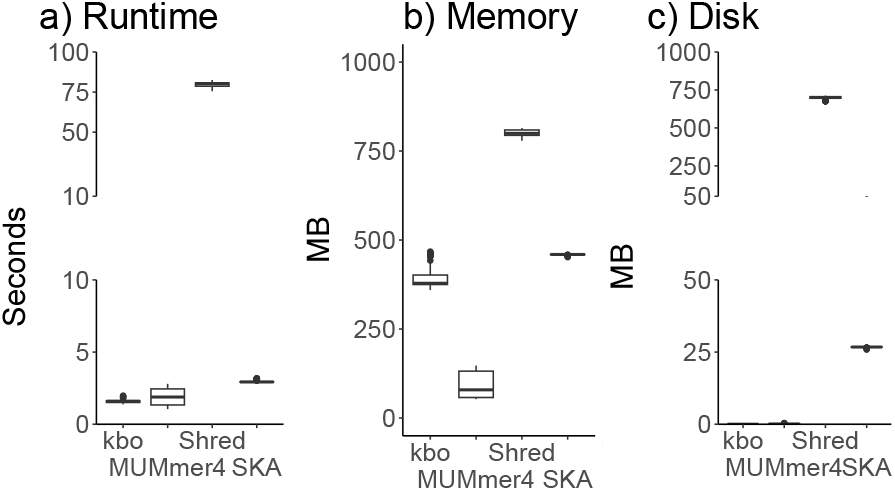
Resource consumption in variant calling for the four tested methods kbo, MUMmer4, Shred, and SKA. Panel a) shows a boxplot of time taken to process each sample. Panel b) the peak memory usage during each run, and Panel c) the temporary disk requirement. The thick lines denote the median and the whiskers the 25% and 75% quantiles. In panels a) and c) the vertical axis has been cut to highlight the differences between Shred and the other methods.

### Gene finding with ‘kbo find’

The ‘find’ command in kbo can be used to find the coordinates of some reference sequence(s) of interest in a query. In this example we demonstrate a use case for ‘find’ in locating 19 genes (total 50,000 bases long) belonging to the pks pathogenicity island in 1999 *E. coli* chromosome sequences (avg. length 4.63Mb) from the Norwegian surveillance programme on resistant microbes (NORM) collection [13, 5].

The ground truth was generated by annotating the 1999 assemblies with bakta [20]. Reference sequences for the pks island genes were extracted from the *E. coli* IHE3034 strain (ASM2574v1A) [17]. Assemblies were processed in four parallel processes with a single thread each using the GNU parallel tool [22], taking a total of 509 seconds (0.25s per assembly) and 195MB peak memory. For comparison, running the same analysis with blast+ [7] takes 364 seconds (0.18s per assembly) and 189MB peak memory. Our tool was able to identify the exact same regions as bakta in 894 out of 934 assemblies. In the remaining 39 assemblies, the average difference between alignments reported by kbo and the gene annotations from bakta was 5 bases out of the total 50 000.

The default mode for ‘find’ does not report the name of the reference sequence (gene or contig, for example) in which an alignment was found due to limitations of the matching statistics algorithm. If the names are required, we implement a detailed mode that indexes each sequence separately, overcoming this problem. The detailed mode is significantly slower, taking a total of 6540 seconds (3.3s per assembly) and 1417MB peak memory to run the same experiment. We leave a more efficient algorithm to future work.

### Reference-based alignment with ‘kbo map’

The final operation supported by kbo is reference-based alignment of two assemblies, where the nucleotides in a query sequence are mapped relative to a reference sequence. The results of this procedure are useful in, for example, phylogenetic inference when the original sequencing reads are not available or the number of queries is prohibitively large for storing the read data. We created a benchmark for ‘kbo map’ by extracting the short-read sequencing data and 112 fully assembled chromosome sequences from *E. coli* sequence type (ST) ST131 isolates in the previously mentioned NORM collection [13, 5]. The ground truth was inferred by selecting a random hybrid assembly as the reference, aligning paired-end short-read sequencing reads from the other isolates against it using BWA-MEM2 [23], and using samtools [8] to convert the alignment into a consensus sequence.

We compared ‘kbo map’, SKA[10], and Snippy[21] to the ground truth by measuring the results (`A, C, G, T`d, and ambiguous or gap) in the alignment that agree or disagree between the tested tools and the ground truth. In the benchmarks, we used assemblies as the inputs to all tools. Our results show that all three tools have comparable performance, with kbo being slightly better when measured by median errors (Tables 3, 4). Figure 7c shows how the results from kbo change when the *k*-mer size and error probability parameters are varied.

**Table 3.** Absolute errors (base pairs) in reference-based alignment of 112 *E. coli* chromosome sequences across kbo, SKA and Snippy.

| Method | 25%-q. | Median | 75%-q. |
| --- | --- | --- | --- |
| kbo | 29 690 | 73 814 | 84 420 |
| SKA | 78 547 | 104 990 | 117 813 |
| Snippy | 124 386 | 149 894 | 159 206 |

**Table 4.** Relative errors (% of reference length) in reference-based alignment of 112 *E. coli* chromosome sequences across kbo, SKA and Snippy. The reference sequence length was 5,025,582 base pairs.

| Method | 25%-q. | Median | 75%-q. |
| --- | --- | --- | --- |
| kbo | 0.6% | 1.5% | 1.7% |
| SKA | 1.6% | 2.1% | 2.4% |
| Snippy | 2.5% | 3.0% | 3.2% |

**Fig. 7:**
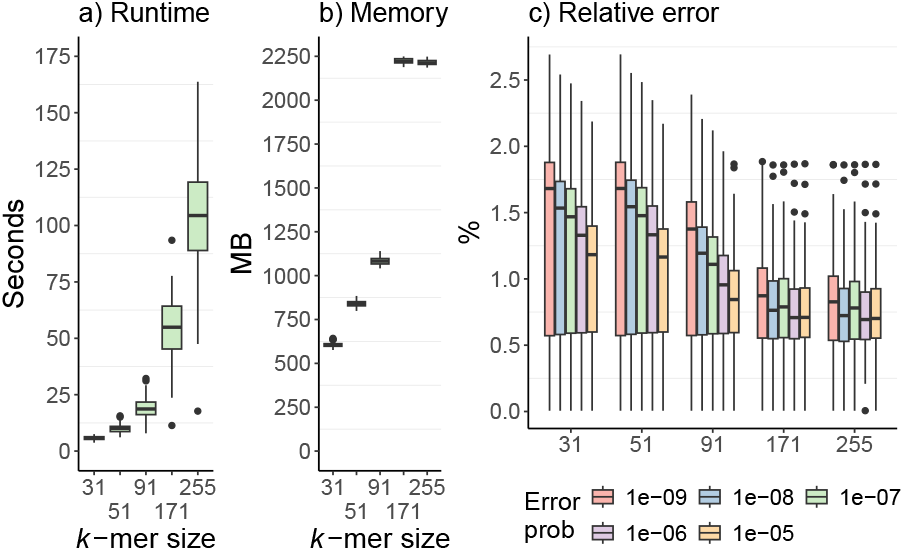
Resource usage and relative error of ‘kbo map’ in reference-based alignment of 112 *E. coli* ST131 chromosome sequences with different parameter configurations. Panel a) shows a boxplot of runtimes of each sequence, Panel b) peak memory usage, and Panel c) relative error measured as number of incorrect base pairs out of the 5 025 582 reference base pairs.

In resource usage kbo and SKA were comparable, with kbo total 624 seconds (mean 5.7s per assembly) and 655MB peak memory to process the benchmark and SKA 669 seconds (6.0s per assembly) and 830MB peak memory. The slowest tool was Snippy which took 2 361 seconds (21.1s per assembly) but had a lower peak memory consumption at 385MB. Additionally, SKA and Snippy used a maximum of 29MB and 52MB temporary disk space, respectively, during the benchmark. The resource usage of kbo is primarily determined by the *k*-mer size parameter (Figures 7a and 7b) and whether further refinmenet with the gap filling and variant calling algorithms are used. Taken together, kbo was the fastest and most accurate method in this benchmark.

### Running kbo in a browser

In addition to the command-line client, we also provide a WebAssembly version of kbo that runs entirely in the browser and does not send any user data to a remote server. The web version of kbo allows users without command-line familiarity to access the tool on any device that with support for a modern browser and modest compute power. Thanks to the frugal resource usage of kbo, the web version supports all functionality provided by the command-line client and can be used to perform any of the analyses presented in this paper. A demonstration of the web version is available online^4^ and can easily be deployed on any web server that supports serving WebAssembly files.

## Discussion

Plain *k*-mer matching methods typically do not support retrieval of the coordinates of the matches. We sidestepped this problem by augmenting *k*-mer matching on the SBWT [3] with *k*- bounded matching statistics [1, 2] and indexing one sequence while streaming the queries from the other, allowing us to transform the matching results into a format where alignment locations can be inferred with high accuracy. Our results show that in addition to being competitive with other methods in accuracy, our approach retains the attractive lookup speed enabled by the specialized data structures for indexing *k*-mers, making it a novel and practically feasible alternative to existing local alignment methods.

One disadvantage of our method is the comparatively high memory usage, especially for large values of *k*, required for constructing the *k*-mer index to perform queries. This issue affects most *k*-mer indexing methods, as the index size is in some way proportional to the number of unique *k*-mers in the input. Index construction algorithms often rely on streaming from temporary disk space to reduce the space usage. We have opted for indexing fully in-memory in order to enable the browser implementation and make our methods better suited for high-performance computing environments, where disk throughput is a limiting factor for parallelisation.

The other future area of improvement is extending matching statistic computation to support coloured index structures, where each *k*-mer in the index is accompanied by a color that denotes its sequence of origin. Storing distinct sequences in this manner would enable efficient indexing of many closely related sequences using color set compression and allow developing kbo towards aligning sequencing reads against large reference collections where it is important to know which targets the read aligned against. The SBWT data structure is easily augmented with color information (witness the Themisto pseudoalignment tool [4]), but an algorithm for colored matching statistics remains an open problem. We leave support for sequencing read alignment and colored matching statistics to future work.

To our knowledge, kbo is the first sequence aligner that compiles to WebAssembly and allows running all features of the tool in a browser including using custom indexes. Web tools have found wide adoption among applied researchers but they remain difficult to customise and require server-side processing, which comes with privacy and long-term support concerns. Using WebAssembly alleviates all of these issues, as the codebase becomes much simpler without a separate client-server model and no data transfer taking place beyond serving the executable to the user. We have made the source code for all parts of our implementation freely available and expect that this will facilitate future implementations of customised analysis pipelines in a manner that is accessible to both non-experts and bioinformaticians alike.

## Code availability

The Rust library, command-line interface, and graphical user interface implementing the methods described here are all freely available from GitHub https://github.com/tmaklin/kbo and Codeberg https://codeberg.org/themaklin/kbo under a MIT and Apache 2.0 dual license. Code for reproducing the benchmarking results is available from Zenodo https://doi.org/10.5281/zenodo.15321495.

## Author contributions statement

T.M. designed the core kbo algorithms and implemented the kbo, kbo-cli, and kbo-gui Rust libraries. J.N.A. implemented the SBWT Rust library and derived the random match distribution. J.N.A. and E.B. designed and implemented the variant calling algorithm. T.M., J.N.A, E.B., and S.J.P. investigated the relationship between *k*-bounded matching statistics and local alignments. S.J.P. obtained funding and supervised the study. All authors contributed to the writing, reviewing, and editing of the manuscript.

## Acknowledgments

This work was supported in part by the Research Council of Finland via grants 339070 and 351150. T.M. was partially funded by Research Council of Norway, grant no. 299941.

## Footnotes

1 https://github.com/tmaklin/kbo

2 https://github.com/tmaklin/kbo-cli

3 https://github.com/tmaklin/kbo-gui

4 https://maklin.fi/kbo

