## Supplementary figures and images for "Sequence alignment with *k*-bounded matching statistics"

### Supplementary Figure 1

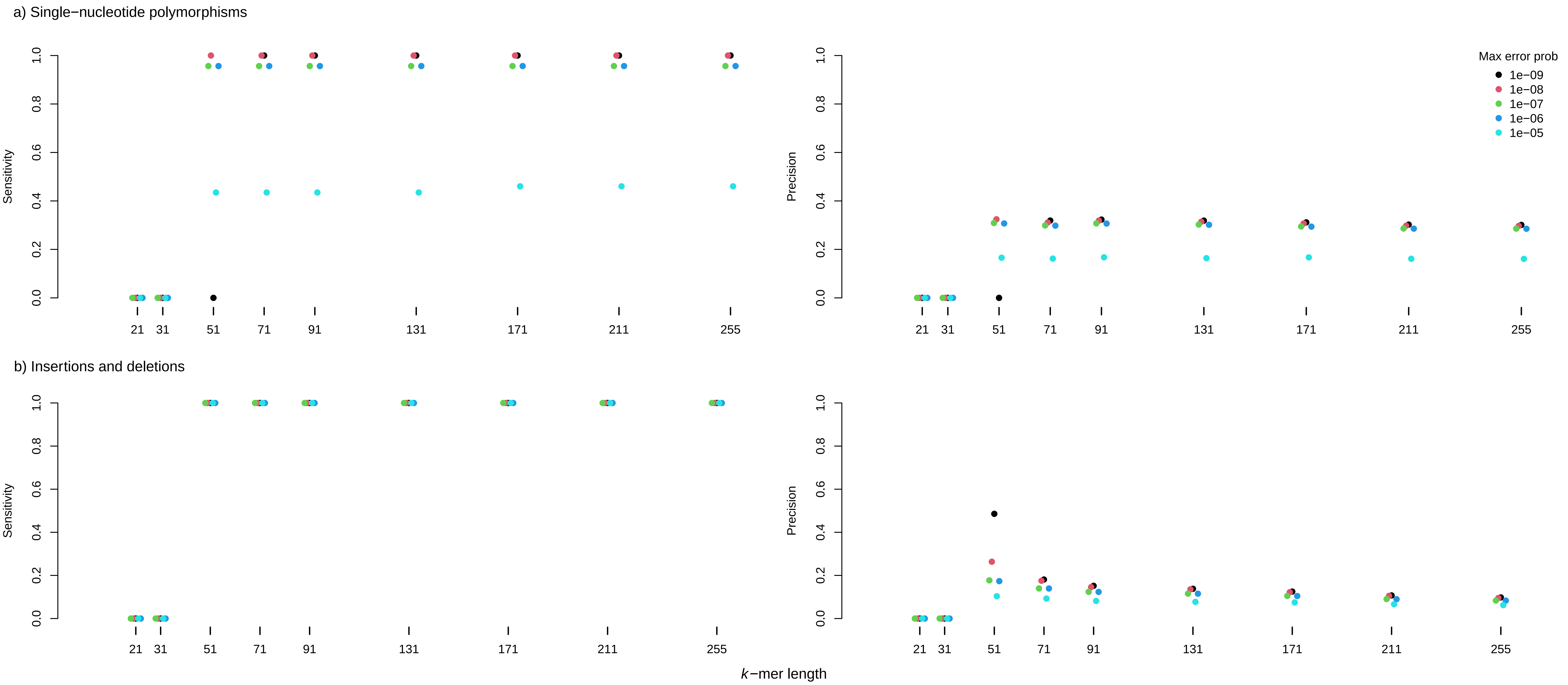

### Supplementary Figure 2

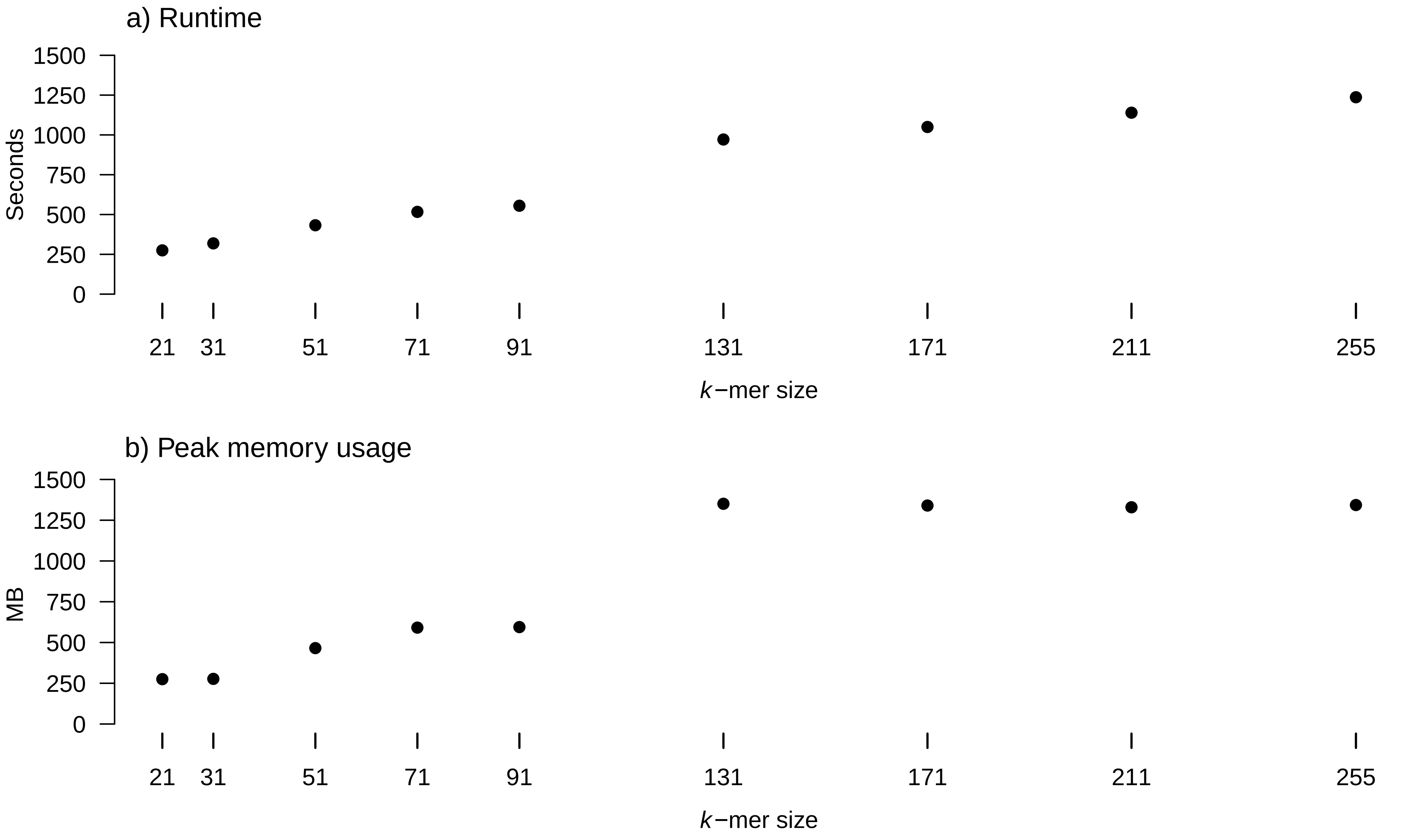
